# A Deep Boosting Based Approach for Capturing the Sequence Binding Preferences of RNA-Binding Proteins from High-Throughput CLIP-Seq Data

**DOI:** 10.1101/086421

**Authors:** Shuya Li, Fanghong Dong, Yuexin Wu, Sai Zhang, Chen Zhang, Xiao Liu, Tao Jiang, Jianyang Zeng

## Abstract

Characterizing the binding behaviors of RNA-binding proteins (RBPs) is important for understanding their functional roles in gene expression regulation. However, current high-throughput experimental methods for identifying RBP targets, such as CLIP-seq and RNAcompete, usually suffer from the false positive and false negative issues. Here, we develop a deep boosting based machine learning approach, called DeBooster, to accurately model the binding sequence preferences and identify the corresponding binding targets of RBPs from CLIP-seq data. Comprehensive validation tests have shown that DeBooster can outperform other state-of-the-art approaches in predicting RBP targets and recover false negatives that are common in current CLIP-seq data. In addition, we have demonstrated several new potential applications of DeBooster in understanding the regulatory functions of RBPs, including the binding effects of the RNA helicase MOV10 on mRNA degradation, the influence of different binding behaviors of the ADAR proteins on RNA editing, as well as the antagonizing effect of RBP binding on miRNA repression. Moreover, DeBooster may provide an effective index to investigate the effect of pathogenic mutations in RBP binding sites, especially those related to splicing events. We expect that DeBooster will be widely applied to analyze large-scale CLIP-seq experimental data and can provide a practically useful tool for novel biological discoveries in understanding the regulatory mechanisms of RBPs.

## 1 Introduction

RNA binding proteins (RBPs) play important roles in multiple aspects of gene expression regulation, such as alternative splicing, RNA modification, mRNA export and localization [1]. Not only does the dysregulation of RBPs induce abnormality, but also the mutations in their binding targets have the potential to cause diseases [2]. So, capturing the intrinsic binding preferences of RBPs and identifying their binding targets in a precise and large-scale manner are essential to understand the regulatory roles of RBPs and reveal their connections to the pathogenesis of complex diseases.

Before the development of high-throughput techniques for characterizing RNA-protein interactions, only a few RBPs were well studied based on the small-scale experiments, such as *in vitro* EMSA [3] and *in vivo* fluorescence methods [4]. Recently, several high-throughput sequencing-based approaches, e.g., CLIP-seq [5–7], SELEX [8,9] and RNAcompete [10,11], have been proposed to measure RBP binding sites and binding affinities in a transcriptome-wide manner. However, despite the huge amount of data generated by these techniques, they still suffer from the false positive and false negative issues mainly due to experimental noise and bias [12]. To overcome these drawbacks, various computational models [13–19] have been developed to learn RBP binding preferences and detect putative RBP targets based on abundant experimental data. As many RBPs have been validated to recognize structured regions [20], there is a tendency in recent studies to incorporate the structural features of target RNAs into prediction models, such as MEMERIS [15], GraphProt [17] and our recent deep learning based model [19], where the integration of RNA structural information has been shown to largely boost the prediction performance. Nevertheless, the current transcriptome-wide experimental techniques for measuring RNA structures are far from maturity. On the other hand, predicting RNA structures using computational models usually requires a substantial amount of additional effort and time, and a predicted RNA structure is generally less accurate compared to that derived from experimental approaches. In addition, systematic integration of both sequence and structural information generally requires a more complex prediction model. So far, it remains largely unknown whether we can derive a sequence based prediction model that only takes RNA seqeuence as input, while still achieving prediction performance comparable to that of the state-of-the-art prediction methods that require both sequence and structural profiles. To fill this gap between modeling accuracy and computational complexity, we develop a deep boosting based model, called DeBooster, that requires only sequence information and can capture RBP binding preferences and predict binding sites from high-throughput CLIP-seq data with high accuracy and efficiency.

Through testing on 24 CLIP-seq datasets, we have shown that even without using RNA structural information, DeBooster can outperform the state-of-the-art methods that take both sequence and structural information as input, including both GraphProt [17] and our previous deep learning based model [19]. In addition, we have performed comprehensive tests to validate the superiority of DeBooster: (i) DeBooster can accurately capture RBP binding preferences and generate RBP binding motifs that are consistent with previous studies in the literature; (ii) DeBooster can successfully carry out the cross-platform prediction task and effectively address the false negative problem that is prevalent in current CLIP-seq data; (iii) In addition to binary classification, DeBooster can be easily extended to solve a regression problem such that the prediction scores closely match the experimentally-measured binding affinities.

In addition to the above extensive validation tests, we have further demonstrated several new possible applications of DeBooster in studying the regulatory roles of RBPs. With an integrative analysis based on other types of data and our prediction results, we not only derive literature-consistent results concerning RBP regulation, but also hope to gain novel insights into the biological rationale of the regulatory roles of RBPs. In particular, we have conformed that the binding targets of the RNA helicase MOV10 predicted by DeBooster are highly associated with the fold changes of mRNA half-lives, providing another evidence on the regulatory functions of RNA helicases on mRNA half-lives. In addition, it has been suggested that a fraction of ADAR binding events might be “non-productive”, i.e., these bindings may not trigger any RNA editing [21]. Consistent with this hypothesis, we have also observed a clear discrepancy in the predicted ADAR binding patterns between “non-productive” and “productive” binding behaviors, which may indicate the existence of different ADAR binding modes to accomplish its diverse regulatory functions. Moreover, we applied DeBooster to study the antagonizing effect of RBP binding on miRNA repression. In particular, it has been known that in the 3 UTR of the oncogene *ERBB2*, the RBP ELAVL1 (also called HUR) antagonizes the repression effect of the miRNA miR-331-3p by binding to a U-rich element (URE) near the miRNA target region called miR-331b [22]. With a mutant URE, we have observed that the new ELAVL1 binding sites predicted by DeBooster shift to a position more distant from the miR-331b region, which is largely consistent with the previous experimental studies. At last, we have used DeBooster to predict the effects of the single nucleotide variant (SNV) mutations on the RBP binding sites related to splicing events, which may provide useful hints for identifying pathogenic mutations and investigating their connections to the pathogenesis of complex diseases. Based on these test results, we expect that DeBooster will have great application potentials and be widely used by the community to analyze more CLIP-seq experimental data and discover more biologically relevant findings on the functional roles of RBPs in post-transcriptional gene regulation.

## 2 Results

### 2.1 The DeBooster framework

We have developed a deep boosting based approach, called DeBooster, to predict the sequence specificities of RNA-binding proteins (RBPs) from high-throughput CLIP-seq data (Fig 1). As RNA primary sequence can be viewed as a string over the alphabet {A, U, C, G}, we mainly use the basic bag-of-words model [23] as in the nature language processing field to encode the features of a given RNA sequence (Fig 1a). In particular, for each word of fixed length *k*, we count how many times it appears in the RNA sequence and store its frequency information in a vector of length 4^*k*^. We extract the word frequency information for both an RBP target region and its upstream and downstream flanking regions of 150 nucleotides each. We consider words of lengths 1, 2, 3, which results in 2 × (4 + 4^2^ + 4^3^) = 168 features in total.

**Figure 1.**
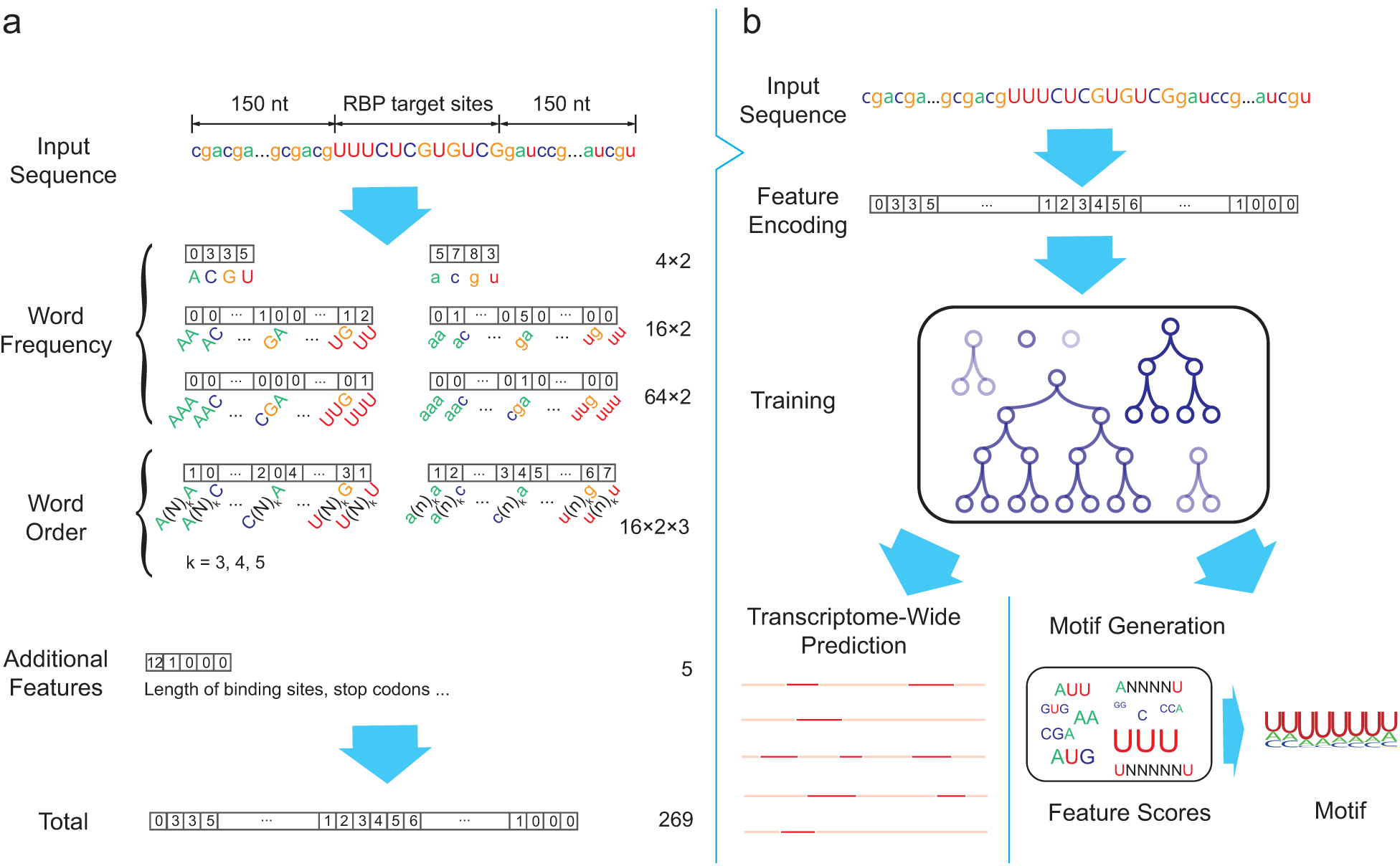
Schematic overview of DeBooster, a deep boosting approach for identifying the sequence specificities of RNA-binding proteins (RBPs). **(a)** Schematic illustration of the strategy for encoding the sequence features of RBP binding targets. The nucleotides in the target region of an input sequence are represented by capitalized letters while the extended regions on both sides are represented by lowercase letters. Each number within a box stands for the value of the corresponding feature. The numbers on the right side represent the total number of feaures in individual catagories. **(b)** Schematic illustration of the prediction pipeline. More details can be found in the main text.

Note that the bag-of-words model mainly focuses on the occurrences of words and reflects little about the order of the letters in a sequence. In other words, if we swap the first half and the second half of an RNA sequence, the features provided by the bag-of-words model would roughly remain the same. To better incorporate the order of letters into the model, we futher use the following scheme to extract the ‘second-order’ word count information. For a fixed stride *m* and a given RNA sequence *a*_1_*a*_2_ … *a_t_*, we count the words *a*_1_*a_m_*_+1_, *a*_2_*a_m_*_+2_, …, *a_t−m_a_t_* and use a vector to record the corresponding count information. As before, we also consider both an RBP target region and the flanking regions of 150 nucleotides both upstream and downstream. We consider the stride lengths 4, 5 and 6, which generates 2 × 3 × 4^2^ = 96 more features in total. Moreover, we consider five additional features, such as the length of the target region, whether the word length is a multiple of 3, whether the target region contains the stop codons UAG, UAA and UGA. Thus, overall we extract 168 + 96 + 5 = 269 features for a given RNA sequence.

We then apply a deep boosting based method, to learn a classification model from the above encoded features (Fig 1b). The deep boosting method [24] is a generalization of several well-established learning approachs, such as AdaBoost [25] and logistic regression [26]. It uses decision tree models as base classifiers and partitions the decision trees of different depths into different sets, denoted by *H*_1_*, …, H_k_*, respectively, where *H_i_* stands for the set of decision trees of depth *k*. In our deep boosting method, we aim to learn a classifier from a family of convex ensembles 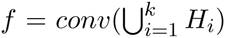. That is, *f* can be written in the form of 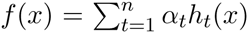, where *α_t_ ≥* 0, and *h*_*t*_ ∈ *H*_*pt*_ for some *p_t_ ∈* [1*, k*]. During the training process, the deep boosting method seeks to minimize the following objective function:

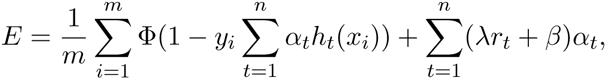

where (*x_i_, y_i_*) denotes the *i*-th training sample, ϕ strands for the total number of training samples, Φ stands for the loss function (e.g., the exponential cost function as in AdaBoost [25] or the logitstic function [26]), *r_t_* stands for the Rademacher complexity of set *H_pt_*, and *λ* and *β* are two hyperparameters to be chosen. The above objective function can be optimized as in other boosting algorithms [25,27]. After training, the learned model can be used to predict the sequence specificities and investigate the corresponding binding motifs of the RBP targets (Fig 1b, Methods).

### 2.2 DeBooster captures the sequence preferences of RBP binding

We first ran a 10-fold cross-validation procedure for each of 24 CLIP-seq datasets (Methods) to evaluate the overall prediction performance of DeBooster. The hyperparameters in the deep boosting framework were determined using an independent dataset (Methods). We also compared the performance of DeBooster with the state-of-the-art approaches for predicting RBP target sites, including GraphProt [17] and the deep belief net (DBN) method [19]. The comparison results (Figs 1a-1c) showed that DeepBooster can significantly outperform both GraphProt and the DBN method, with the increase of the area under receiver operator characteristic curve (AUROC) by up to 10.1%. Note that GraphProt and the DBN method integrate both RNA sequence and structural information (i.e., RNA secondary structural information [17] or both RNA secondary and tertiary structural profiles [19]) into the prediction framework, while DeBooster requires only RNA sequence information. The performance improvement in DeBooster was probably attributed to our new feature encoding scheme (see Fig 1a and Section 2.1) and the better predictive power of the underlying deep boosting model.

Through a transcriptome-wide analysis on RBP binding targets, we also found that the difference in the predicted binding scores of DeBooster over different characterized genomic regions mostly reflected the known functions of individual RBPs (Supplementary Notes). In addition, we examined the sequence motifs of the RBP binding sites generated from training data (Methods). Our results indicated that the sequence motifs resulting from DeBooster agreed well with those reported in the literature (Fig 2d). For example, the binding sequence motif of AGO2 computed by DeBooster was enriched with A, U and C but depleted of G, which was consistent with the previous study [28]. PTB, as indicated by its name (polypyrimidine tract-binding protein), mainly binds to the U/C-rich regions [29], which was also reflected in the sequence motif derived from DeBooster. EWSR1, FUS and TAF15 belong to the FET family. Although several works showed that they bind to the GU-rich motif [30,31], recent studies found that the FET protein family prefers binding to the AU-rich stem loops, and the AU-rich sequences achieve higher binding affinities than those enriched with G and U [32]. Such an AU-rich pattern was also observed in the sequence motif generated by DeBooster. It has been found that the binding targets of QKI usually contain a core sequence NACUAAY (where Y stands for a pyrimidine) and a half-site UAAY [33]. The biniding motif of QKI identified by DeBooster also agreed well with such a pattern. DeBooster yielded a U-rich sequence motif for the binding sites of HNRNPC, which can also be supported by a known fact that HNRNPC generally binds to the poly-U tracts [34]. According to the DeBooster prediction results, SFRS1 prefers binding to a GA-rich motif, which aligned well with the previous result [35]. As shown in the previous study [7], PUM2 binds to a consensus motif UGUANAUA, which shared high similarity with the corresponding binding motif predicted by DeBooster. The majority of the TDP43 binding sites predicted by DeBooster contained the (UG)_*n*_ motif and was relatively less enriched with A and C. Such an observation agreed well with the previous known result [36]. Taken together, most of the sequence motifs of RBP binding sites captured by DeBooster were consistent with the previous known results in the literature.

**Figure 2.**
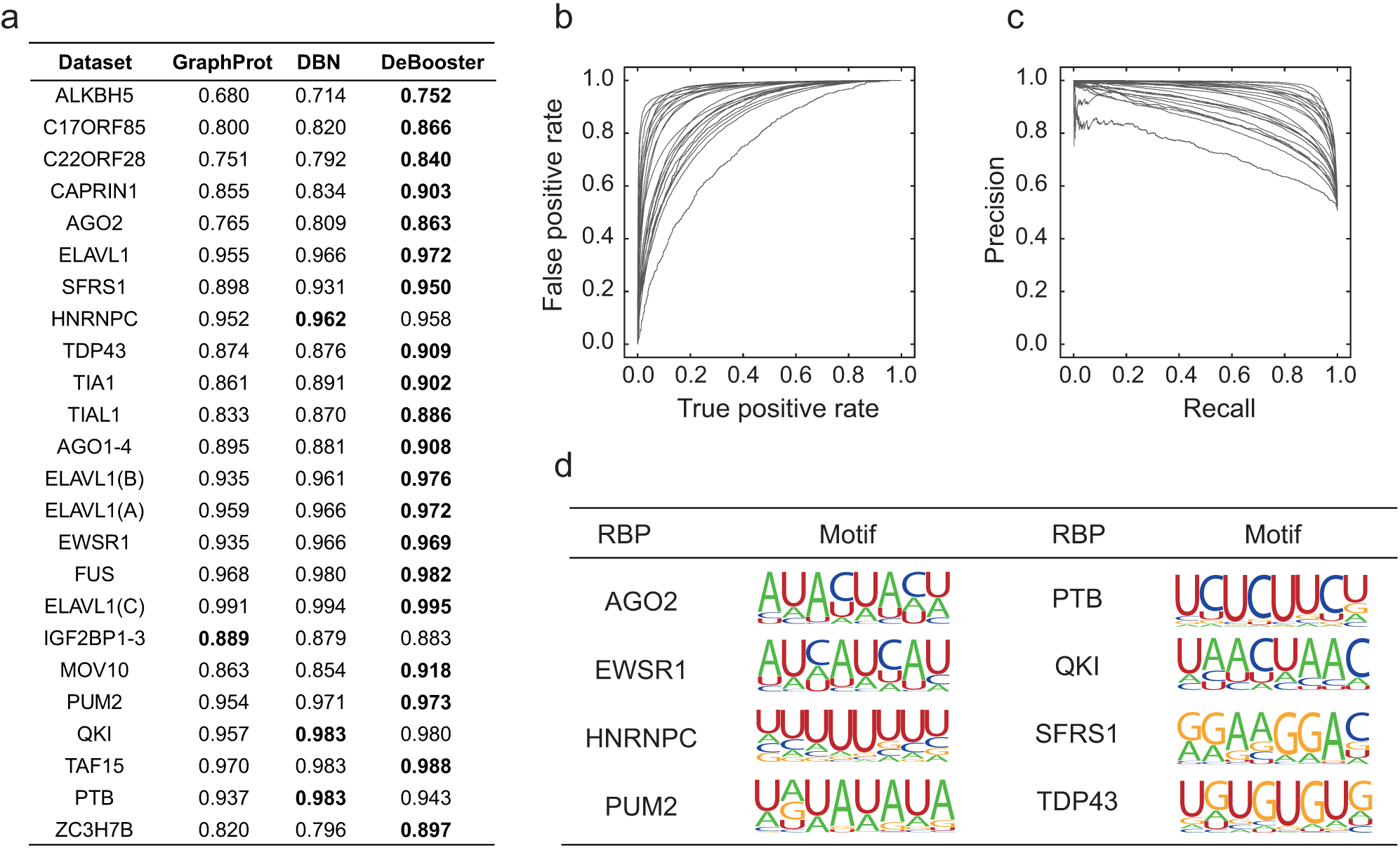
Performance evaluation of DeBooster on 24 CLIP-seq datasets. **(a)** The comparisons of the area under receiver operator characteristic curve (AUROC) scores between different prediction approaches via a 10-fold cross-validation procedure. The best prediction result for each dataset is highlighted in bold. **(b)** and **(c)** The receiver operator characteristic (ROC) and precision-recall (PR) curves achieved by DeBooster for all 24 CLIP-seq datasets in the cross-validation results, respectively.

### 2.3 The predictions of DeBooster can be validated through cross-platform datasets

It is well-known that different CLIP-seq experiments can yeild a large fraction of non-overlapping results and individual experiments may miss a vast number of true RBP binding sites [37,38]. Here, we showed that the prediction results of DeBooster can be validated through cross-platform datasets and thus effectively alleviate the false negative problem in current high-throughput CLIP-seq data (Fig 3). In particular, we tested DeBooster on different cross-platform ELAVL1 datasets, which displayed a large degree of discrepancy between the original RBP binding targets measured from CLIP-seq experiments (Fig 3a). Such a large variation indicated that in general a single CLIP-seq experiment cannot cover all RBP binding sites and individual datasets may have high false negative rates in current experimental measurement. The tests on the cross-platform ELAVL1 datasets showed that the predictions of DeBooster from one dataset can be well validated by another one collected from a different platform, achieving both high AUROC scores and similar sequence motifs (Fig 3b). In addition, most of the sequence features encoded in DeBooster displayed highly correlated weights except the outliers G and UNNNNU (Figs 3c-3d), which was probably due to experimental bias introduced from the original CLIP-seq data. Thses results implied that the predictions of DeBooster can be well validated through cross-platform CLIP-seq datasets.

**Figure 3.**
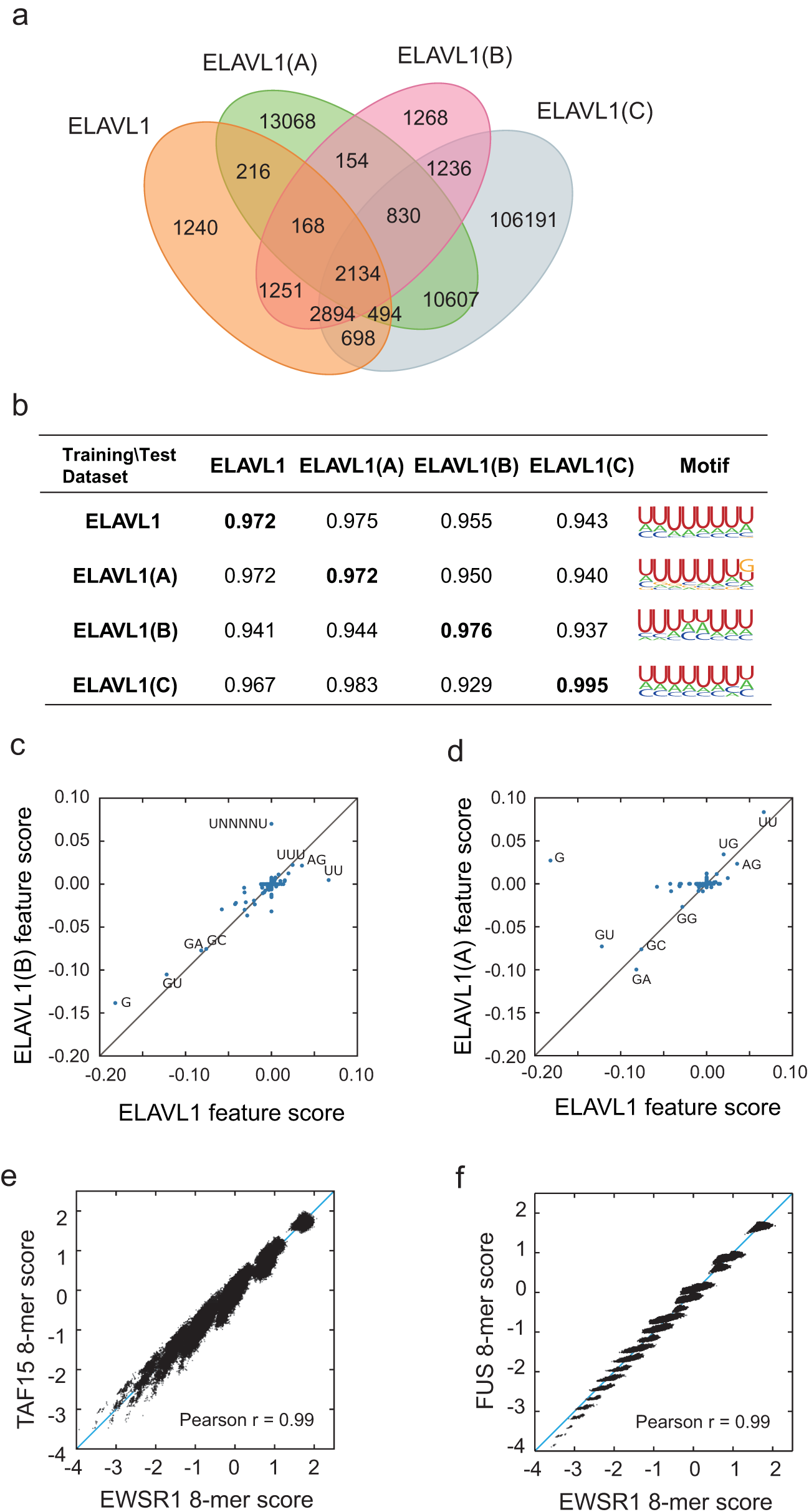
Performance validation of DeBooster through cross-platform CLIP-seq datasets. **(a)** The Venn diagram of four ELAVL1 CLIP-seq datasets collected from different experimental platforms. If binding region A from a dataset has at least one nucleotide overlap with binding region B from another dataset, we regarded A and B as a common element of these two datasets. The datasets ELAVL1, ELAVL1(A) and ELAVL1(C) were from the HEK293 cells, while the dataset ELAVL1(B) was from the Hela cells. **(b)** The AUROC scores and binding sequence motifs computed by DeBooster using different combinations of training and test datasets. The diagonal scores shown in bold correspond to the cross-validation results in which both training and test datasets were collected from the same experimental platform. **(c, d)** The plots of the relative weights of individual sequence features computed by DeBooster for the ELAVL1 datasets collected from different experimental platforms, including ELAVL1(B) vs. ELAVL1 **(c)** and ELAVL1(A) vs. ELAVL1 **(d)**. **(e, f)** The plots of the DeBooster prediction scores for all 8-mers across different RBPs within the same family, including TAF15 vs. EWSR1 **(e)** and FUS vs. EWRS1 **(f)**. TAF15, FUS and EWSR1 all belong to the FET family and generally share similar binding preferences.

We also investigated the agreement of the DeBooster prediction results between different RBPs from the same family. In particular, we examined the consistency between the DeBooster prediction scores of 8-mers for TAF15, FUS and EWSR1, all belonging to the FET family. Consistent with the previous results that these three RBPs have a large overlap in binding sites [32], our tests showed that the 8-mers from different RBPs exhibited highly correlated prediction scores (Figs 3e-3f). Such observations further supported the above argument that the prediction results of DeBooster can be verified from cross-platform CLIP-seq datasets, even for different RBPs from the same family. These results suggested that DeBooster was not prone to overfitting, and may provide a practically useful tool to analyze high-throughput CLIP-seq data and recover false negatives that are common in current CLIP-seq data.

### 2.4 The binding scores predicted by DeBooster match the experimentally measured binding affinity data

To investigate whether the prediction results of DeBooster can truely reflect the RBP binding preferences, we further checked the agreement between the binding scores predicted by DeBooster and the experimentally determined binding affinity data. In particular, we compared the predicted binding scores with both *in vivo* determined K_d_ values [39,40] and *in vitro* measured binding affinities from RNAcompete assays [10,11].

We first checked the agreement between the prediction scores of DeBooster, which was trained using the *in vivo* CLIP-seq data, and the experimentally determined K_d_ values for two RBPs, including SFRS1 and TDP43 (Figs 4a-4b). Our comparison showed that for the 8-mers as the potential RNA targets of SFRS1, the prediction scores of DeBooster closely matched the *in vivo* measured K_d_ values [39] (Fig 4a). In addition, for the RNA nucleotides as the potential binding targets of TDP43, the prediction scores of DeBooster aligned well with the K_d_ values experimentally measured from the electrophoretic mobility shift assay (EMSA) [40] (Fig 4b).

**Figure 4.**
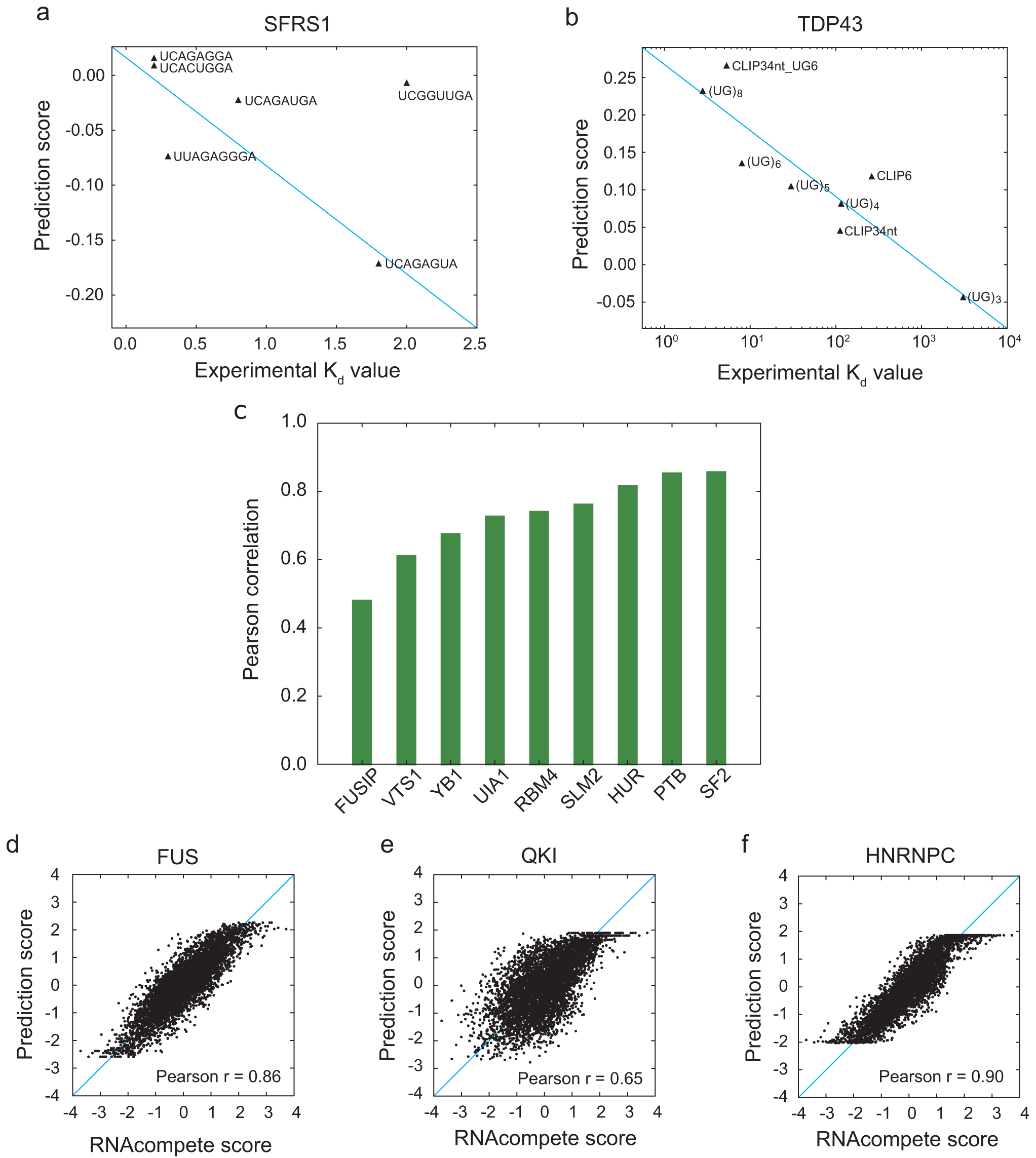
The comparisons between the prediction scores derived by DeBooster and the experimentally determined binding affinity data. **(a, b)** The plots of the prediction scores derived by DeBooster (which was trained based on CLIP-seq data) vs. the experimentally determined K_d_ values of different 8-mers or RNA oligonucleotides for SFRS1 and TDP43, respectively. The K_d_ values of SFRS1 were measured *in vivo* [39], while the K_d_ scores of TDP43 were acquired from the electrophoretic mobility shift assay (EMSA) [40]. The same terminology as in [40] for the names of RNA oligonucleotides was used for the binding targets of TDP43 (Supplementary Notes). **(c)** The Pearson correlation coefficients between the prediction scores derived by DeBooster vs. the *in vitro* binding affinity scores of 7-mers derived from the RNAcompete data in [10]. **(d-f)** The plots of the prediction scores derived by DeBooster vs. the *in vitro* exprimentally measured binding affinity scores of 7-mers derived from the RNAcompete data in [11] for FUS, QKI and HNRNPC, respectively. In **(c-f)**, an extended “regression” version of DeBooster was used and a cross-validation procedure was applied to evaluate the prediction performance (Methods).

Next, we examined the prediction performance of DeBooster on the *in vitro* binding data derived from the RNAcompete experiments [10,11]. The original version of DeBooster took binary (i.e., positive or negative) samples as input. To enable the model to consider the real-valued RNAcompete data, we also made a simple extention and proposed a “regression” version of DeBooster (Methods). A cross-validation test (Methods) showed that the prediction scores of DeBooster and the *in vitro* binding scores from the RNAcompete assays [10,11] for the synthesized oligonucleotides displayed high correlations. In particular, the comparisons to the earlier version of the RNAcompete data [10] for the synthesized 7-mers of nine RBPs exhibited high consistency, in which most of Pearson correlation efficients were above 0.7 (Fig 4c). In addition, the comparisons to the latest version of the RNAcompete data [11] for three RBPs, including FUS, QKI and HNRNPC, showed good agreement between the prediction scores and the *in vitro* binding affinity scores, with Pearson correlation coefficients above 0.65 (Figs 4d-4f). Overall, these comparison results implied that the binding scores predicted by DeBooster may be regarded as a useful indicator of RBP binding affinities for both *in vivo* and *in vitro* scenarios.

### 2.5 The predicted targets of RNA helicases may be associated with the regulation of mRNA degradation

RNA helicases, such as MOV10, regulate the life cycle of mRNAs and thus gene expression by remodeling RNA secondary structures and RNA-protein interactions [41]. Here, we showed that the RNA targets of MOV10 predicted by DeBooster can be connected to the regulation of mRNA half-lives and thus may provide useful hints for understanding the funcitonal roles of MOV10 in controlling gene expression. Our analysis was performed on a set of 7000 mRNAs, in which the fold changes of their half-lives had been measured after MOV10 knockdown [42]. These mRNAs were basically divided into four groups according to the fold changes of their half-lives, i.e., top 25%, 25%-50%, 50%-75% and bottom 25%, which corresponded to Group 1, Group 2, Group 3 and Group 4, respectively. Only the bottom group (i.e., Group 4) contained those genes whose expression levels were unchanged or up-regulated after MOV10 knockdown.

Compared to the results derived directly from the original CLIP-seq data (Fig 5a), the fraction of UTRs with MOV10 binding resulting from DeBooster prediction displayed a more evident decreasing trend (Fig 5b). In addition, the sum of all positive prediction scores per UTR, which basically considered both binding strength and the number of hits for the MOV10 binding targets on individual genes, also exhibited the same decreasing order for four groups of genes that were divided and ranked according to the fold changes of mRNA half-lives (Fig 5c). Moreover, when we grouped all transcripts according to the DeBooster prediction scores, the resulting fold changes of mRNA half-lives also presented a similar decreasing trend (Fig 5d). Furthermore, the DeBooster predition scores for seven genes also showed good agreement with the fold changes of mRNA half-lives experimentally measured by qRT-PCR (Fig 5e). Taken together, the above results demonstrated that the binding targets of the RNA helicase MOV10 predicted by DeBooster were associated with the changes of mRNA half-lives. Thus, the prediction results from DeBooster may provide useful clues for further understanding the regulatory machanisms of RNA helicases on the life cycle of mRNAs.

**Figure 5.**
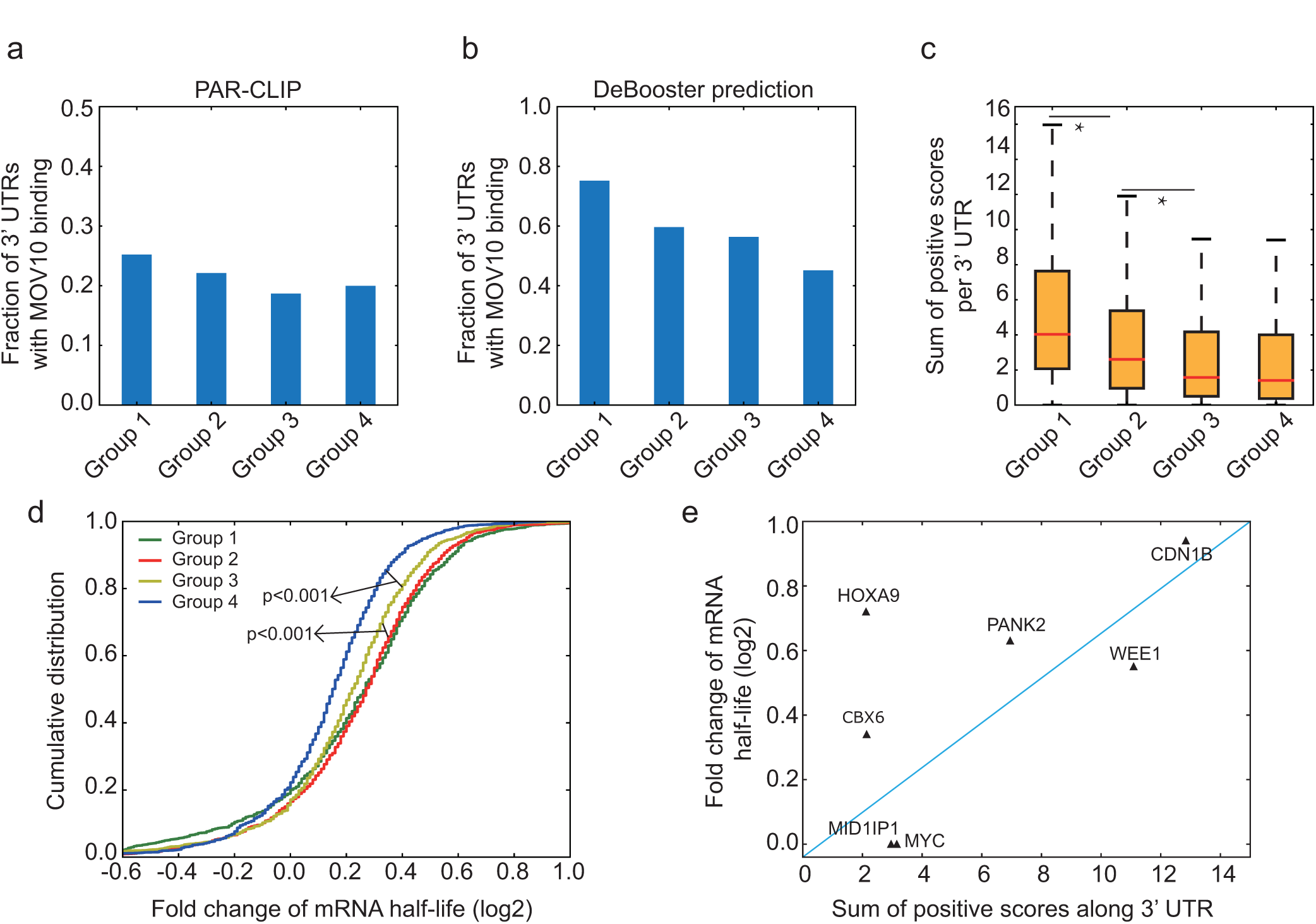
Understanding the predicted binding effects of MOV10 on mRNA dagradation. **(a, b)** Fractions of 3 UTRs with MOV10 binding for four groups classified according to the original CLIP-seq data **(a)** and the DeBooster prediction results **(b)**, respectively. Genes were evenly separated into four groups according to the fold changes of their mRNA half-lives. Groups 1, 2, 3 and 4 corresponded to top 25%, 25%-50%, 50%-75% and bottom 25%, respectively. In the DeBooster prediction results, we only considered those robust binding targets with prediction scores *>* 0.2 (The default threshold was zero and the range of prediction scores was in [-1,1]). **(c)** The sum of positive prediction scores per UTR for four groups of genes, which were classified and ranked according to the fold changes of their mRNA half-lives in a descending order. *: p value *<* 0.001, Wilicoxon rank sum test. **(d)** The cumulative distribution on the fold changes of mRNA half-lives for four groups of genes, classified and ranked according to the DeBooster prediction scores in a descending order. That is, Groups 1, 2, 3 and 4 corresponded to genes with top 25%, 25%-50%, 50%-75% and bottom 25% predicted scores, respectively. The p values were computed using the Wilicoxon rank sum test. **(e)** The plot of the DeBooster prediction scores vs. the fold changes of mRNA half-lives meansured by qRT-PCR for seven genes.

### 2.6 Applying DeBooster to study the difference between productive and nonproductive ADAR binding patterns

ADARs are a family of homologous enzymes catalyzing adenosine-to-inosine (A-to-I) editing in the RNA, and have similar double-stranded RNA binding domains (dsRBDs) and a common deaminase domain [43]. Despite their major role as RNA-editing enzymes, a fraction of ADAR binding events might be “non-productive”, that is, these bindings might not trigger any RNA editing [21]. On the contary, those ADAR binding events that indeed produce RNA editing were considered “productive”. To investigate the difference of the binding behaviors between productive and non-productive ADAR binding targets, we compared the prediction results from three DeBooster models, which were trained using all, productive and non-productive ADAR binding sites, respectively.

We first introduced the concept of the binding-editing distance, which was defined as the genomic distance between an ADAR binding position and its closest editing site. The known RNA editing sites were obtained from the RADAR database [44]. Our first model, also called the all-binding model, was trained using all ADAR1 binding sites measured from CLIP-seq experiments [45] as the postive samples. The negative samples were defined as those unbound regions that were adjacent to the positive samples in transcripts and had the lengths equal to those of the corresponding positve samples. In our second model, also called the productive binding model, the CLIP-seq sites (i.e., the ADAR1 binding sites measured from CLIP-seq experiments) with small binding-editing distances (0−100 nt) were used as the positive samples, while the CLIP-seq sites with large binding-editing distances (>1000 nt) together with the adjacent unbound regions were used as the negative samples. In our third model, also called the non-productive binding model, the CLIP-seq sites with large binding-editing distances (>1000 nt) were used as the positive samples, while the CLIP-seq sites with small binding-editing distances (0−100 nt) together with the adjacent unbounded regions were used as the negative samples. The median of the binding-editing distances resulting from the all-binding model was 814 nt (Fig 6a), which was roughly on the same scale as from the original CLIP-seq data (606 nt). The median of the binding-editing distances from the productive binding model was zero (i.e., the ADAR binding region contained at least one editing site), which was significantly different from that of the non-productive binding model (4665 nt, Fig 6a). The above results implied that indeed DeBooster may be able to distinguish productive and non-productive ADAR binding sites based on current high-throughput CLIP-seq data.

**Figure 6.**
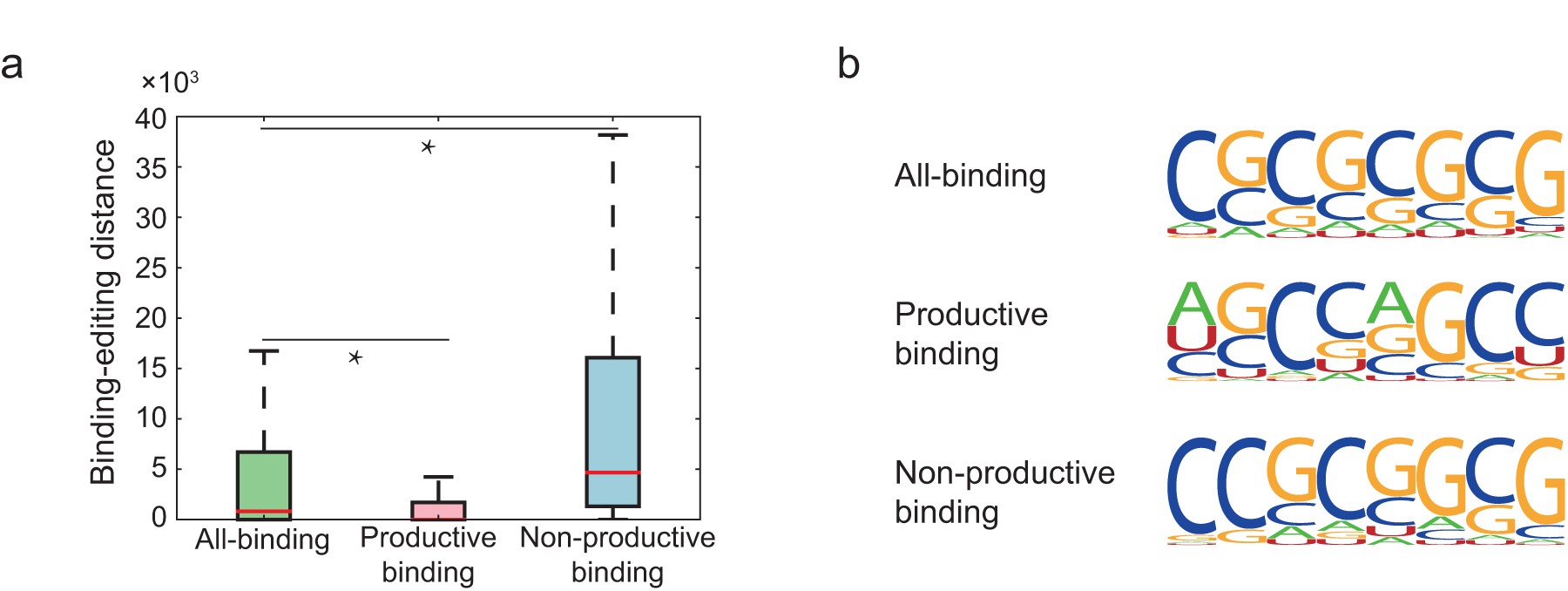
The comparison results on three different DeBooster models, which were trained using all ADAR binding sites identified by CLIP-seq experiments, productive ADAR binding sites (i.e, triggering A-to-I editting), and non-productive ADAR binding sites (i.e, without triggering A-to-I editting), respectively. **(a)** The boxplot of the binding-editing distances, which were defined as the genomic distances between the ADAR binding sites and the closet editing sites, for three different DeBooster models. *: p value *<* 0.001, Wilicoxon rank sum test. **(b)** The sequence motifs of the ADAR binding sites identified by three different DeBooster models. More details can be found in the main text.

We also examined the sequence motifs of the ADAR binding sites identified by three different DeBooster models (Fig 6b). Although all three sequence motifs showed high GC content, the motif generated by the productive-binding model had relatively higher frequencies of As and Us than those from the other two models. This observation indicated that those ADAR binding sites with relatively lower GC content might be more prone to being edited. This result was also in agreement with the known evidence that the published motif of the ADAR binding sites [45] contained relatively higher GC content than that of the genomic regions near the editing sites [46].

As in previous studies [21], our results showed that some ADAR binding sites were close to the editing sites, while many others were thousands of nucleotides away from the editing sites. Although it was possible that this phenomenon was attributed to the lack of the complete editing records in the database, the clear discrepancy of the binding-editing distances and the sequence motifs of the ADAR binding sites between productive and non-productive models derived from DeBooster indicated that there may exist different binding modes to accomplish diverse regulatory effects of ADAR binding.

### 2.7 The shift of the predicted RBP binding scores from mutations may predict the antagonizing effect of RBP binding on miRNA repression

RBPs and miRNAs are two classes of essential regulators controlling mRNA degradation and expression, and they often interplay with each other to display co-regulatory effects [47]. For example, in the 3' UTR of an oncogene *ERBB2,* the RBP ELAVL1 (also called HUR) antagonizes the repression effect of the miRNA miR-331-3p by binding to a U-rich element (URE) near the miRNA target region called miR-331b [22]. With a mutant URE, the repression effect of ELAVL1 binding on miR-331-3p is weakened, since the new ELAVL1 binding sites shift to a position that is more distant from the miR-331b region (Fig 7a), and also reduces the binding affinity of ELAVL1 (the magnitude of the experimentally measured K_d_ values change from 10^−8^ M to 10^−7^ M) [22]. Here, we showed that DeBooster can successfully identify this mutational effect that was consistent with the previous experimental observation.

**Figure 7.**
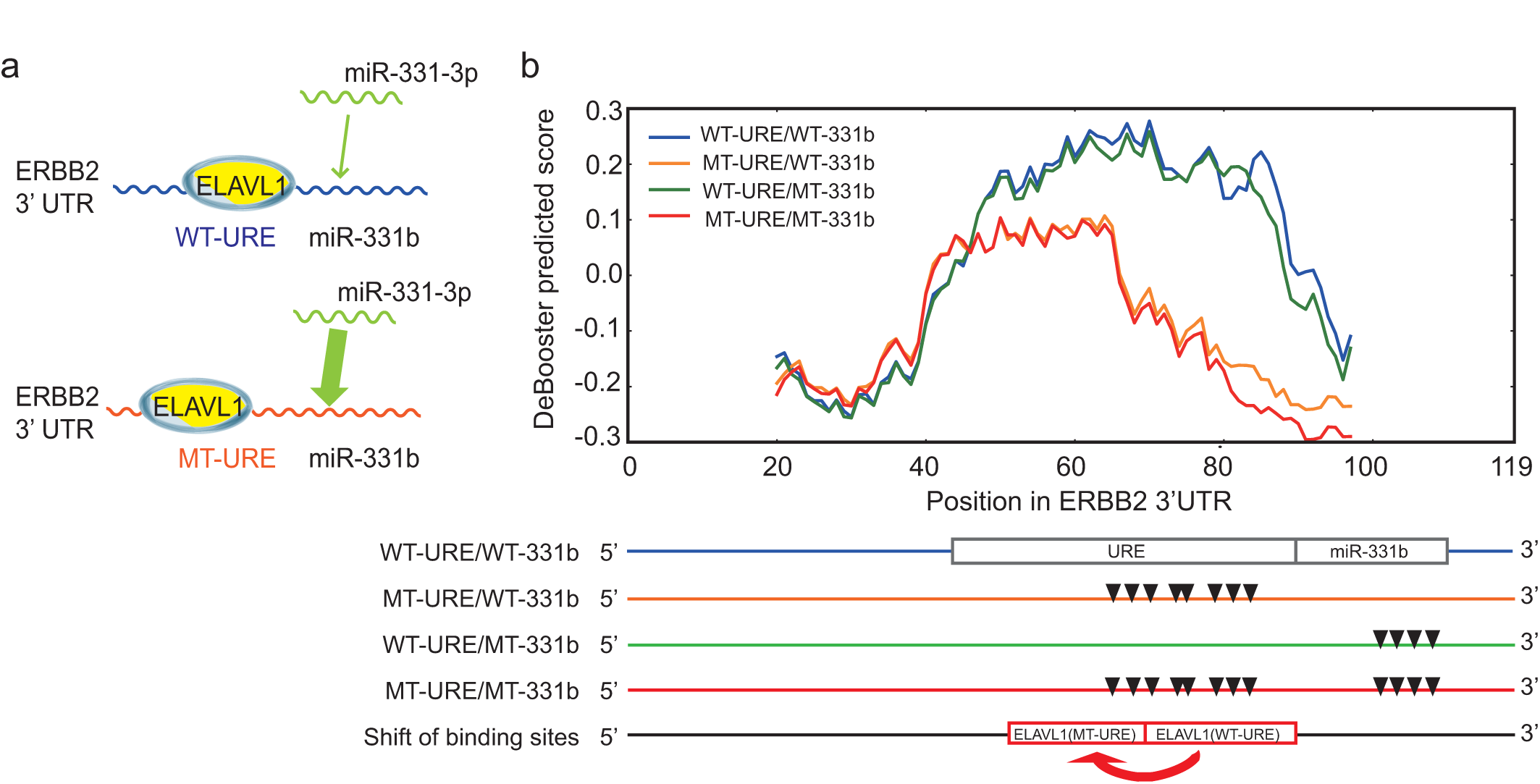
The predicted influence of ELAVL1 binding on the repression effect of miRNA miR-331-3p. **(a)** An illustrative model of the co-binding of RBP ELAVL1 and miRNA miR-331-3p on the 3 UTR of gene *ERBB2.* miR-331b represents the binding region of miRNA miR-331-3p. The width of the arrow represents the relative strength of miR-331-3p binding. **(b)** The change of the predicted binding scores corresponded to the shift of ELAVL1 binding sites from the wild-type to the URE mutant on the 3 UTR of gene *ERBB2.* The bottom shows the locations of URE and miR-331b regions, mutation positions in the URE region, mutation positions in the miR-331b region, mutation positions in both URE and miR-331b regions, and the experimentally detected shift of ELAVL1 binding resulting from the URE mutant, respectively. All mutation sites are represented by the inverted triangles. Abbreviation: WT, wild-type; MT, mutant; URE, U-rich element.

We used the CLIP-seq dataset of ELAVL1 measured from the Hela cells [48] as training data (those overlapping records about the measured binding sites in the 3' UTR of gene *ERBB2* were removed) and performed a comparative study on the predicted binding scores of four cases, i.e., WT-URE/WT-331b, MT-URE/WT-331b, WT-URE/MT-331b and MT-URE/MT-331b, which represented the wild-type sequence, a URE mutant with the wild-type miR-331b region, the wild-type URE with a miR-331b mutant, and a sequence with mutations in both URE and miR-331b regions, respectively (Fig 7b). All the binding scores predicted by DeBooster showed obvious peaks near the URE, indicating the high-affinity binding of ELAVL1 in this region. More importantly, the prediction results of DeBooster displayed a clear position-shifted and affinity-decreased pattern of ELAVL1 binding on a URE mutant (Fig 7b). The curves of the predicted binding scores for WT-URE/WT-331b (i.e., wild-type) and WT-URE/MT-331b (i.e., only mutations in the miR-331b region) had similar shapes, which was consistent with the previous experimental result that the mutations in the miR-331b region rarely affect ELAVL1 binding [22]. In addition, the peaks of these two curves were approximately located in positions 50-90 along the 3' UTR of ERBB2, while the peaks of the other two curves with mutations in the URE region (i.e., MT-URE/WT-331b and MT-URE/MT-331b) were located around positions 45-60. Such a position shift of the ELAVL1 binding sites identified by DeBooster in fact agreed with the previous experimental RNA footprinting results (see Fig 7b in [22]). Morever, the decrease of the binding scores predicted by DeBooster was also consistent with the loss of the experimentally-determined K_d_ values with respect to the same mutations [22]. Taken together, these results indicated that DeBooster can successfully identify the changes of the RBP binding scores caused by the mutations in binding targets which may be used to predict the antagonizing effect of RBP binding on miRNA repression.

### 2.8 The prediction scores of DeBooster may provide a useful index to study pathogenic mutations affecting RNA splicing

Recent studies revealed that abnormal splicing play a vital role in development of many human diseases, such as cancer and neurological disorders [49–51]. The mutations near splice sites or on splicing regulatory elements, such as exonic splicing enhancers (ESEs) and exonic splicing silencers (ESSs), may influence RNA splicing and cause human diseases by disrupting RBP binding [2]. Here, we were particularly interested in whether DeBooster can identify pathogenic mutations and thus be used as a useful tool to study the mutational effects of sequence variants related to splicing events. We first examined the overall changes of the predicted binding scores of individual RBPs with respect to the sequence variants of their binding targets near 5' and 3' splice sites (Methods) and checked whether DeBooster was able to distinguish pathogenic mutations from neutral sequence variants. Our comparisons showed that the changes of the binding scores predicted by DeBooster for a majority of pathogenic sequence variants in regions near 5' and 3' splice sites were significantly different from those of neutral mutations (Fig 8). In addition, almost all of these pathogenic mutations displayed relatively larger changes in the predicted binding scores than neutral variants. On the other hand, most of the neutral mutations near 5' and 3' splice sites displayed similar effects (with only 4 among 20 RBPs showing significant difference with p*<*0.001 in the Student’s t test). Furthermore, the pathogenic mutations near splice sites generally showed a greater extent of difference in the predicted binding scores than those pathogenic mutations randomly chosen from the COSMIC records [52] (Supplementary Fig 2). For instance, among 20 RBPs, 15 and 18 proteins exhibited significantly different mutational effects on the pathogenic variants near 5' and 3' splice sites, respectively, compared to only 7 RBPs in those pathogenic mutations randomly selected from COSMIC (Fig 8 and Supplementary Fig 1). Such an observation implied that the seqeuence disruptions of the RBP binding targets around splice sites may generally play a more important role in the pathogenisis of a disease. Overall, our studies indicated that the binding scores derived from DeBooster may provide an effective indicator for distinguishing pathogenic mutations from neutral variants in RBP binding targets near splice sites.

**Figure 8.**
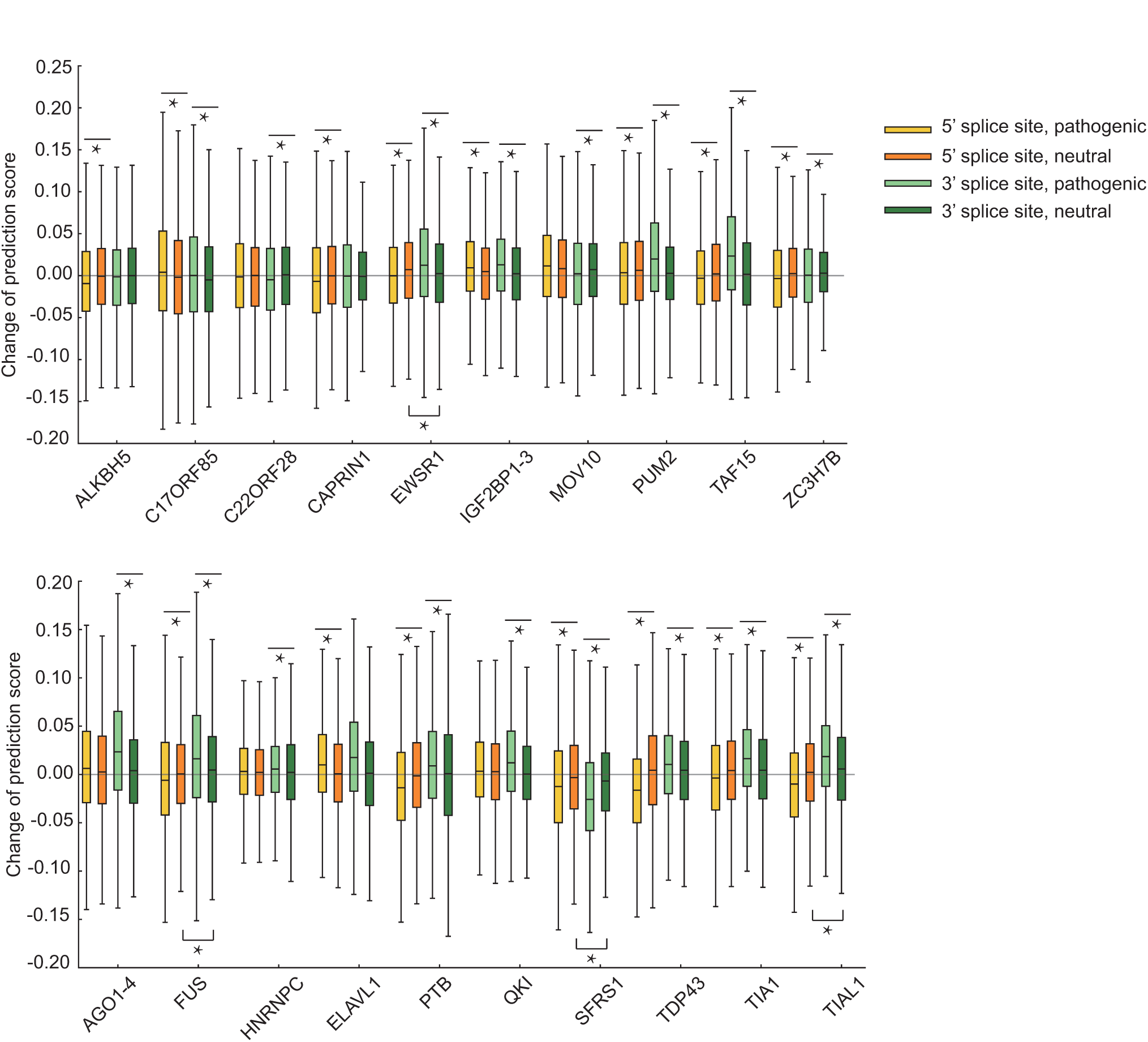
The comparisons between the overall changes of the predicted binding scores of individual RBPs after pathogenic or neutral mutations in regions near 5 and 3 splice sites. *: p*<*0.001, Student’s t test.

Next, we further analyzed the mutational effects predicted by DeBooster for a number of known pathogenic single-nucleotide variants (SNVs) obtained from COSMIC [52]. Below we describe several examples (Fig 9). First, a synonymous substitution of the last base in Exon 7 (G to A) of gene *CDH1* (which encodes the E-cadherin protein) led to an increase in the SFRS1 binding scores predicted by DeBooster near a 5 splice site (Fig 9a), which may be related to the dysregulation of *CDH1* that causes tumor metastasis [53]. Such an observation may also be supported by a previous experimental validation study that this mutation can actually alter splicing by causing intron retention to various extents [54].

**Figure 9.**
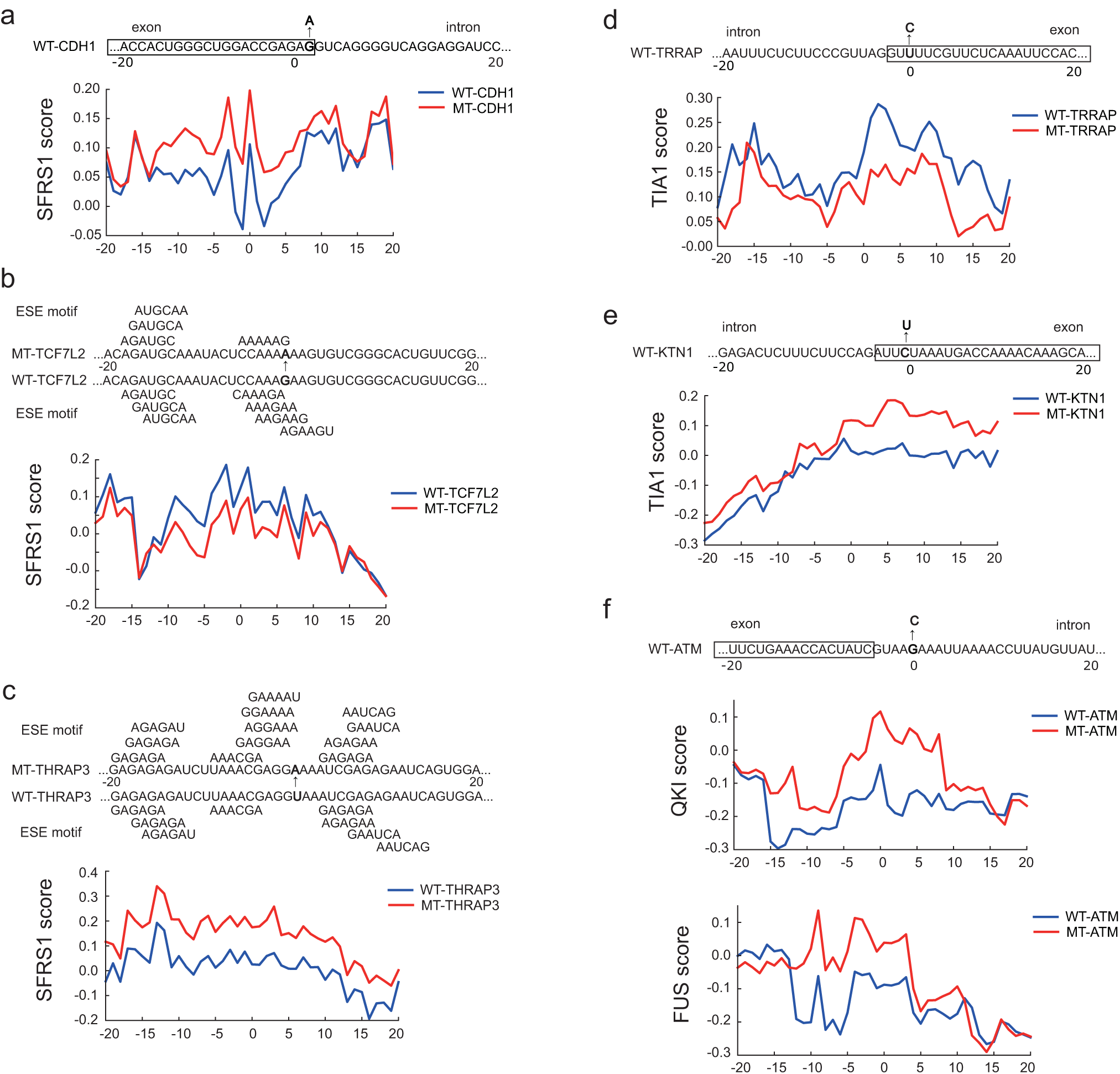
Examples of the predicted effects on the potentially disease-causing mutations near splice sites or on exonic splicing enhancers (ESEs). **(a)** The exonic mutations of the SFRS1 binding sites near a 5 splice site for gene *CDH1*. **(b, c)** The mutations of SFRS1 binding sites disrupting or creating exonic splicing enhancer (ESE) motifs for genes *TCFIL2* and *THRAP3*, repectively. The ESE motifs were obtained from [65]. **(d, e)** The exonic mutations of the TIA1 binding sites near the splice sites for genes *TRRAP* and *KTN1*, respectively. **(f)** A mutation near a 5 splice site of gene *ATM* that changed the predicted binding scores of both QKI and FUS. Abbreviation: WT, wild-type; MT, mutant; URE, U-rich element.

As a second example, a mutation from G to A in a *TCF7L2* exon [55] disrupted the ESE motifs (which are 6 nt motifs located in exons and bound by SR proteins to promote exon splicing [56]) and suppressed SFRS1 binding (Fig 9b), while a mutation from U to A in a *THRAP3* exon [55] enriched the ESE motifs and thus enhanced SFRS1 binding (Fig 9c). Such disruptions in those disease-relevant genes may influence the binding behaviors of the important splicing regulator SFRS1, and thus may be related to the tumorgenesis associated with aberrant splicing [39].

In our third example, the mutation from U to C near a 3 splice site of gene *TRRAP* [55] weakened TIA1 binding (Fig 9d). TRRAP interacts with oncoproteins MYC and E2A [57], and its mis-regulation can be heavily related to various types of cancers [58]. On the other hand, another mutation from C to U near a 3 splice site of gene *KTN1* [55] strengthened TIA1 binding (Fig 9e). *KTN1* encodes kinectin 1, and has been shown to display different splicing patterns in cancers [59]. Thus, these two sequence variants in the binding sites of TIA1 may be associated with cancer pathogenesis by changing the alternative splicing modes of its target genes.

Another interesting example is the intronic mutation near a 5 splice site of gene *ATM* [55], which increased the binding scores of both FUS and QKI (Fig 9f). Such a mutation may influence the splicing result of this tumor suppressor (i.e., ATM) [60] by creating new potential binding sites for both splicing regulators (i.e., FUS and QKI).

In addition to the above cases, there were other examples to demonstrate that the prediction scores of DeBooster may reflect the pathogenic effects of sequence disruptions in RBP binding. For instance, a substitution from C to U near a 5 splice site of gene *NF1* [55] enhanced HNRNPC binding (Supplementary Fig 3a), which may be associated with the known related neurologic disorders [61]. On the other hand, a mutation from U to C near a splice site of the proto-oncogene *BRAF* [55] decreased the HNRNPC binding score (Supplementary Fig 3b). In addition, a mutation from A to G [55] near a splice site of gene *TET2* may help form a novel GU-repeat region for strong TDP43 binding (Supplementary Fig 3c), and thus influence the splicing process. Moreover, the *SMAD4* splicing site may be disrupted by the mutation from G to U [55] that may increase the PTB binding score (Supplementary Fig 3d) and thus alter the corresponding splicing result. Both *TET2* and the *SMAD4* genes act as tumor suppressors [62,63], so the inhibition of their normal splicing may thus facilitate cancer formation.

Taken together, the above examples illustrated that the RBP binding scores predicted by DeBooster may offer a useful index to investigate the pathogenic effects of sequence disruptions related to RNA splicing.

## 3 Conclusion

We developed DeBooster, a deep boosting based framework to model the sequence binding specificities of RNA-binding proteins (RBPs) from high-throughput CLIP-seq data. Compared to the state-of-the-art methods which usually require both sequence and structure profiles, DeBooster uses only sequence information as input. Tests on 24 CLIP-seq datasets demonstrated that DeBooster can achieve better prediction performance than previous methods. Through a validation test on several cross-platform CLIP-seq datasets of ELAVL1, we showed that DeBooster can be useful for addressing the false negative problem that is prevalent in current CLIP-seq data. In addition, the prediction scores of DeBooster agreed with the experimentally-determined binding affinity scores, such as *in vivo* measured K_d_ values and the *in vitro* binding affinities measured from RNAcompete.

We further showed the great application potentials of DeBooster by applying it to study the regulatory roles of several important RBPs. In particular, we demonstrated that the predicted targets of the RNA helicase MOV10 can better explain its binding effects on the regulation of mRNA degredation than the original CLIP-seq data. In addition, the predicted RBP binding sites may help understand the difference between productive and non-productive binding patterns of the RNA-editing enzymes ADAR. We also showed that a shift of the predicted ELAVL1 binding scores from wild-type to mutant in a U-rich element (URE) region of gene *ERBB2* can effectively predict the antagonizing effect of RBP binding on miRNA regulation. Moreover, DeBooster may be used as an effective index to identify pathogenic mutations from normal sequence variants and study the effects of potential disease-causing mutations in RBP binding sites related to splicing. Based on these test results and analyses, we expect that DeBooster will provide a promising tool to analyze more large-scale CLIP-seq data and gain more biological insights related to RBP regulation.

## 4 Methods

### 4.1 Datasets

We used 24 sets of CLIP-seq based data about RBP binding sites to train and validate our prediction model. These datasets were preprocessed in [17] to generate both positive and negative samples. The list of all RBP names in these 24 CLIP-seq datasets can also be found in Fig 2a. Among these datasets, AGO1-3 and IGF2BP1-3 contained the binding targets of several RBPs from the same protein family, while ELAVL1 HITS-CLIP, ELAVL1 PAR-CLIP(A), ELAVL1 PAR-CLIP(B) and ELAVL1 PAR-CLIP(C) included the binding sites of RBP ELAVL1 measured from different experimental platforms.

### 4.2 Determination of hyperparameters

We use an independent validation dataset of RBP C22ORF28 to determine the optimal setting of the hyperparameters of DeBooster, including the type of the loss function (denoted by Φ), the number of the base decision tree classifiers (denoted by *n*), the maximun depth of these decision trees (denoted by *k*), and parameters *λ*, *β* controlling the relative importance of the complexity penalty. This process yields the following optimal setting of the hyperparameters: the exponential function as the loss function Φ, *n* = 200, *k* = 5, *λ* = 0.3 and *β* = 0.

### 4.3 Motif generation

We use the following procedure to generate representive motifs of the RBP binding sites predicted by DeBooster. First, we use the set of the weighted decision trees resulted from the deep boosting algorithm to evaluate the relative importance of each encoded feature. In particular, for each decision tree with weight *ω* in the model, we identify the feature *ψ* and the corresponding threshold *τ* used to split the root node for this feature. Suppose that at the root node a fraction *p*_1_ of all examples in the training set are postive, and at the right child of the root node (in which the value of feature *ψ* is larger than *τ*), a proportion *p*_2_ of all examples in the training set are positive. We then use (*p*_1_ *− p*_2_)*ω* to represent the importance of feature *ψ*. By doing so, we score each feature based on its contribution to RBP binding. A higher absolute value of a positive score means higher contribution to RBP binding, while a higher absolute value of a negative score means less contribution to RBP binding. We use a vector *s* to store the importance scores of all encoded features. Next, we go through all 8-mers and extract the feature vector *v_i_* for each of them. We then rank these 8-mers according to the inner product of *v_i_* and *s*, and we select the top 500 8-mers with the highest ranking scores. As the top 8-mers may come from shifts around the best one, we align all 8-mers with respect to the top one such that the largest number of base matchings is achieved. After that, we generate the binding motif based on this alignment step and visualize it using the WebLogo site [64].

### 4.4 An extension of DeBooster to real-value labeled data

Although the original version of DeBooster is designed to take only binary (i.e., positive or negative) labeled examples in training data, its output value is nevertheless real-valued, which offers us the possibility to adapt it to handle a regression problem.

To be specific, suppose the training set *S* for the regression task is {(*x*_1_*, y*_1_)*, …,* (*x_n_, y_n_*)}, where each target value (i.e., the label) *y_i_* is real-valued. We now construct a new training set 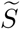 as follows: for each (*x_i_*, *y_i_*) ε *S*, we add an example 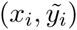 into 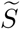, where 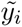 takes value 1 with probability *p_i_*, −1 with probability 1 *− p_i_* and *p_i_* = exp(2*y_i_*)*/*(1 + exp(2*y_i_*)).

In this way, the new training set 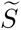 only consists of examples with binary labels (i.e., either 1 or −1), and thus is suitable for classification. In addition, we assert that the ground truth value *y_i_* should minimize the loss in expectation. Therefore, ideally the output value predicted by the model for *x_i_* after training using dataset 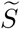 should be equal to *y_i_*.

To see that, recall that our model uses the exponential function as the loss function Φ. Hence, if the original deep boosting model outputs *t_i_* for the *i^th^* example, the expected loss is given by:

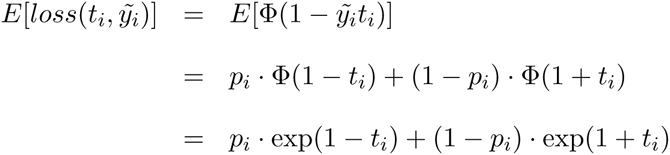

We seek a variable *t_i_* that minimizes the above expected loss by setting the derivative to 0, i.e.,

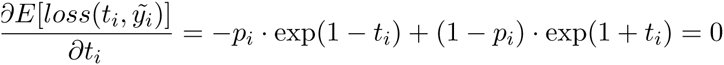

Plugging in *p_i_* = exp(2*y_i_*)*/*(1+exp(2*y_i_*)), we can see that *t_i_* = *y_i_* satisfies the above equation, which means that *t_i_* = *y_i_* minimizes the expected loss. Therefore, in an ideal case the model will output *y_i_* on an input *x_i_* after training based on 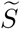.

We also consider several practical issues for the above regression version of DeBooster. First, in order to handle different training sets whose distributions of the target values *y*’s may be largely different, we first preprocess the *y*’s and transform them into a Gaussian distribution. To do so, all *y*’s are sorted and the *i^th^ y* is then transformed into (2 ∗ *i* − 1)*/*(2*n*), where *n* is the number of training examples. In this way, these transformed *y*’s will have a uniform distribution in [0, 1]. Hence it can be further transformed into a Gaussian distribution in a standard way.

Besides, as the output range of the deep boosting model is only within [−1, 1], it would only make sense to guarantee the transformed *y*’s to mainly fall in this range. Therefore, we choose to transform the *y* into a Gaussian distribution with zero mean and standard deviation of 0.4. Notice that 1*/*0.4 = 2.5, which means that we treat those target values *y*’s that are outside 2.5 standard deviations as outliers and force them into [−1, 1] by assigning them with value 1 or −1.

To evaluate the performance of the regression, we first split the dataset into 70% for training and 30% for testing. A transformation function *f* is first learned from the training set, and then used to predict the value *t* for each example in the test set. After that, the Pearson coefficient between *t*’s and and *f*(*y*)’s is used to evaluate the regression performance.

### 4.5 Predicting ELAVL1 binding scores along the 3

Both wild-type and mutant 3' UTR sequences were obtained from [22] (Supplementary Notes). The lengths of these sequences are all 119 nt. For each sequence, we took a window of length 41 nt (the average length of the ELAVL1 target regions over training samples) and slided this window along the 3' UTR of mRNA ERBB2 with a stride length of 1 nt. For each sliding window, we assigned the resulting prediction score to the central nucleotide of this window. Overall, we obtained the prediction scores along positions 21-99 for each sequence (Fig 7), and the first and last 20 nucleotides were not included in our analysis.

### 4.6 Studying the effects of mutations in RBP binding targets

The mutation data related to splicing events were derived from COSMIC [52]. Sequences with mutation sites in the middle and lengths equal to those of the corresponding RBP binding targets were prepared as input samples to DeBooster. For both pathogenic or neutral mutations near 5' or 3' splice sites, we selected those single-nucleotide variant (SNV) mutations within 10 nt from splice sites. The lengths of RBP binding targets are usually larger than 20 nt, so generally splice sites were covered by samples centered at mutation positions. In total, we collected 7,000 neutral mutations in both regions near 5' and 3' splice sites, and 4,000 and 20,000 mutations in regions near 5' and 3' splice sites, respectively. In Fig 8, the change of the prediction score resulting from a mutation was calculated as “(prediction score for the mutant sequence)−(prediction score for the wild-type sequence)”.

In Fig 9 and Supplementary Fig 3, the prediction scores for regions around the mutation sites along both wild-type and mutant sequences were shown. For each selected mutation, we showed the prediction scores for 41 positions, including the mutation site and the flanking regions of 20 nucleotides both upstream and downstream. For each site, its prediction score was calculated using the window centered at this position and of length equal to the average length of the corresponding RBP targets in the training data.

## Acknowlegements

This work was supported in part by the National Basic Research Program of China Grant 2011CBA00300 and 2011CBA00301, the National Natural Science Foundation of China Grant 61033001, 61361136003 and 61472205, US National Science Foundation grants DBI-1262107 and IIS-1646333, China’s Youth 1000-Talent Program and the Beijing Advanced Innovation Center for Structural Biology. The authors are grateful to Dr. Qiangfeng Zhang and Mr. Hailin Hu, Mr. Bin Zhou, and Mr. Xuan He for their helpful discussions about this work.

## Author contributions

S.L., F.D, Y.W. and J.Z. conceived the research project. J.Z. supervised the research project. F.D. and Y.W. designed and implemented DeBooster, and carried out model training and validation tasks. S.L., F.D., S.Z., C.Z., T.J., X.L. and J.Z. performed the computational and statistical analysis. S.L., F.D. and J.Z. wrote the manuscript. All the authors discussed the test results and commented on the manuscript.

